# Aryl Hydrocarbon Receptor Activation Drives 2-Methoxy Estradiol Secretion in Human Trophoblast Stem Cell Development

**DOI:** 10.1101/2024.08.27.609205

**Authors:** Vinay Shukla, Khursheed Iqbal, Hiroaki Okae, Takahiro Arima, Michael J. Soares

## Abstract

**STUDY QUESTION:** How does activation of AHR signaling affect human trophoblast cell development and differentiation?

**SUMMARY ANSWER:** AHR activation leads to altered gene expression but does not hinder the ability of trophoblast cells to remain in a stem cell state or differentiate into essential cell types, such as extravillous trophoblast cells (EVT) or syncytiotrophoblast (ST). It also promotes the production of 2 methoxy estradiol (2ME), a compound that could influence placental development.

**WHAT IS KNOWN ALREADY:** The placenta serves both as a nutrient delivery system and a protective barrier against environmental toxins. AHR signaling is known to mediate cellular responses to environmental pollutants, potentially affecting trophoblast cell functions, but the specific impacts of AHR activation on these cells were not fully understood.

**STUDY DESIGN, SIZE, DURATION:** This study utilized an in vitro model of human trophoblast stem (TS) cells to investigate the downstream effects of AHR activation. The study focused on both undifferentiated TS cells and cells undergoing differentiation.

**PARTICIPANTS/MATERIALS, SETTING, METHODS:** Human trophoblast stem (TS) cells were used as the model system. Researchers examined the effects of TCDD exposure in both TS cells maintained in their stem state and those induced to differentiate into EVT or ST. The study assessed changes in gene expression, particularly focusing on *CYP1A1* and *CYP1B1*, as well as the production of 2ME.

**MAIN RESULTS AND THE ROLE OF CHANCE:** AHR activation stimulated the expression of *CYP1A1* and *CYP1B1*, key genes associated with AHR signaling, in both undifferentiated and differentiating trophoblast cells. While AHR activation did not impact the cells ability to remain in a stem state or differentiate, it increased the production of 2ME, which may influence placental function. These effects were dependent on AHR signaling.

**LIMITATIONS, REASONS FOR CAUTION:** This study was conducted in vitro, which may not fully replicate human conditions. Further research is needed to confirm whether these findings apply to actual placental development in humans.

**WIDER IMPLICATIONS OF THE FINDINGS:** The results suggest that AHR signaling activated by environmental pollutants could have a subtle but significant impact on placental development through mechanisms involving AHR activation. These findings may have broader implications for understanding how environmental factors affect fetal development.

**STUDY FUNDING/COMPETING INTEREST(S):** This work was funded by the National Institutes of Health: ES028957, HD020676, ES029280, HD105734 and the Sosland Foundation. The authors declare no conflicts of interest.

## INTRODUCTION

The placenta is a specialized organ that enables a safe and supportive environment for the fetus to develop within the female reproductive tract. Functional properties of the placenta are attributed to specialized lineages of trophoblast cells (**Soares et al. 2018; Knofler et al. 2019; Shukla et al. 2024**). Disruptions in trophoblast cell differentiation and placental morphogenesis affect fetal development and contribute to the origins of adult disease (**Burton et al. 2016**). There is a myriad of environmental exposures that could impact placentation and embryonic development (**Mattison 2010; Marsit 2016; Vrooman and Bartolomei 2016**). An environmental exposure may lead to placental dysmorphogenesis and dysfunction and/or may exacerbate placental dysfunction in pregnancy-associated diseases (**Gingrich et al. 2020; Zhao et al. 2024**). Timing of environmental exposures is likely critical in determining their effects on placentation and postnatal health (**Barouki et al. 2012**). The impact of environmental exposures on placental development has received limited experimental attention.

Some environmental toxicants affect cellular function through physical interactions with the aryl hydrocarbon receptor (**AHR**) (**Beisclag et al. 2008; McIntosh et al. 2010; Avilla et al. 2020**). These compounds are effectively ligands for AHR and include halogenated aromatic hydrocarbons (e.g. polychlorinated biphenyls, polychlorinated dibenzodioxins, and dibenzofurans), polycyclic aromatic hydrocarbons (e.g. benzo[a]pyrene and benzanthracene), indoles, flavones, benzoflavones, imidazoles, pyridines, lipids, and lipid metabolites (**Birnbaum 1994; Denison and Nagy 2003; Baba and Nakanishi 2005; DeGroot et al. 2012; Murray and Perdue 2020**). AHR is a ligand-activated transcription factor and member of the PER–ARNT– SIM subgroup of the basic helix-loop-helix superfamily of transcription factors (**Vazquez-Rivera et al. 2021**). Upon ligand binding, AHR translocates to the nucleus and heterodimerizes with AHR nuclear translocator (**ARNT**) (**Beisclag et al. 2008; McIntosh et al. 2010**). This heterodimer binds to aryl hydrocarbon response elements (**AHREs**) located within regulatory regions of target genes, including those encoding proteins that are important in biotransformation, drug metabolism, and detoxification of environmental pollutants (**Beisclag et al. 2008; McIntosh et al. 2010; Avilla et al. 2020**). Cytochrome P450 family 1 subfamily A member 1 (**CYP1A1**) is a prototypical transcriptionally activated gene induced by AHR signaling (**Whitlock 1999; Ma 2001**). AHR has been implicated as a regulator of a wide range of biological processes critical for embryonic development and homeostasis as reviewed by Puga and colleagues (**Zablon et al. 2021**).

The barrier for progress in understanding the impact of environmental exposures on placental development is the implementation of appropriate experimental models to test relevant hypotheses. In vitro approaches are powerful. There are wide range of immortalized and transformed cell models that have been used with the goal of elucidating human trophoblast cell responses to AHR ligands (**Zhang et al. 1995, 1997, 1998; Stejskalova et al. 2011, 2013; Tsang et al. 2012; Fadiel et al. 2013; Le Vee et al. 2014; Wu et al. 2016; Dral et al. 2019**).

Unfortunately, deciphering trophoblast cell biology using immortalized and transformed cell models is inherently confounding with questionable relevance (**Lee et al. 2016**). The isolation and culture of trophoblast stem (**TS**) cells from several species, including rodents and primates, represented a major advance for investigating trophoblast cell lineage development (**Tanaka et al. 1998; Asanoma et al. 2011; Okae et al. 2018; Matsumoto et al. 2020; Schmidt et al. 2020**).

In this proposal, we investigated an environmental exposure, 2,3,7,8-tetrachlorodibenzo-p-dioxin (**TCDD**) on trophoblast cell development using human TS cells. TCDD is a well-characterized prototypical activator of AHR signaling (**Wilson and Safe 1998; Mandal 2005; Bock 2018**) and its actions have been investigated in a range of developmental systems (**Couture et al. 1990; Carney et al. 2006; Yoshioka and Tohyama 2019; Yongkai et al. 2024**). We show that TCDD-mediated AHR activation modulates the developmental fate of human TS cells.

## MATERIALS AND METHODS

### Chemicals

2,3,7,8-tetrachlorodibenzo-p-dioxin (**TCDD**, D-404S) was obtained from AccuStandard and solubilized in dimethyl sulfoxide (**DMSO**, D8418, Sigma-Aldrich). 17β estradiol was purchased from Sigma-Aldrich (3301) and solubilized in ethanol.

### Human TS Cell Culture

Cytotrophoblast-derived human TS cell lines (CT27, 46, X,X; CT29, 46, X,Y) were maintained in the stem state or differentiated into extravillous trophoblast (**EVT**) cells or syncytiotrophoblast (**ST**), as described previously (**Okae et al. 2018**). Human TS cells were routinely cultured in 100 mm tissue culture dishes coated with 5 μg/mL of mouse collagen IV (35623, Discovery Labware) or human collagen IV (5022, Advanced Biomatrix). Complete TS Cell Medium was used to maintain cells in the stem state and consisted of Basal TS Cell Medium [DMEM/F12 (11320033, Thermo Fisher), 100 μm 2-mercaptoethanol, 0.2% (vol/vol) fetal bovine serum (**FBS**), 50 μM penicillin, 50 U/mL streptomycin, 0.3% bovine serum albumin (**BSA**, BP9704100, Thermo Fisher), 1% insulin-transferrin-selenium-ethanolamine solution (vol/vol, Thermo-Fisher)] with the addition of 200 μM L-ascorbic acid (A8960, Sigma-Aldrich), 50 ng/mL of epidermal growth factor (**EGF**, E9644, Sigma-Aldrich), 2 μM CHIR99021 (04-0004, Reprocell), 0.5 μM A83-01 (04-0014, Reprocell), 1 μM SB431542 (04-0010, Reprocell), 0.8 mM valproic acid (P4543, Sigma-Aldrich), and 5 μM Y27632 (04-0012-02, Reprocell).

Human TS cells were investigated in the stem state and following differentiation into EVT cells or ST. In these experiments, we utilized a range of TCDD concentrations (1 to 100 nM) to empirically determine concentrations sufficient to induce activation of AHR signaling in vitro.

### EVT cell differentiation

To promote EVT cell differentiation, human TS cells were cultured in 6-well plates pre-coated with 1 μg/mL of collagen IV at a density of 80,000 cells per well.

Cells were cultured in EVT Cell Differentiation Medium, which consists of the Basal TS Cell Medium with the addition of 100 ng/mL of neuregulin 1 (**NRG1**, 5218SC, Cell Signaling), 7.5 μM A83-01, 2.5 μM Y27632, 4% KnockOut Serum Replacement (**KSR**, 10828028, Thermo Fisher), and 2% Matrigel (CB-40234, Thermo Fisher) (**Okae et al. 2018**). On day 3 of differentiation, the medium was replaced with EVT Differentiation Medium excluding NRG1 and with a reduced Matrigel concentration of 0.5%. On culture day 6 of EVT cell differentiation, the medium was replaced with EVT Differentiation Medium excluding NRG1 and KSR, and with a Matrigel concentration of 0.5%. Cells were analyzed on day 8 of EVT cell differentiation.

### ST differentiation

To promote ST differentiation, TS cells were cultured in 6-well plates at a density of 300,000 cells per well using ST-Three Dimensional (**ST3D**) Medium, which consists of Basal TS Cell Medium with a decreased concentration of BSA (0.15%) and the addition of 200 μM L-ascorbic acid, 5% KSR, 2.5 μM Y27632), 2 μM forskolin (F6886, Sigma-Aldrich), and 50 ng/mL of EGF (**Okae et al. 2018**). On day 3 of cell differentiation, 3 mL of fresh ST3D Medium was added to the wells. Cells were analyzed on day 6 of ST differentiation.

### Flow cytometry assay for cell death

Cells (2 × 10^5^ cells/ml) were cultured in 6-well plates and exposed to TCDD (10 nM and 100 nM) for 24 h. Cells were trypsinized, washed with PBS and incubated with FITC-conjugated Annexin-V (catalog number and Vendor?) and probidium iodide (200 µl of a 50 mg/L solution; catalog number and Vendor; for 15 min. Staining profiles were assessed by flow cytometry.

### Cell cycle analysis

Cells (2 × 10^5^ cells/ml) were cultured in 6-well plates and exposed with TCDD (10 nM and 100 nM) for 24 h. Cells were trypsinized, washed with PBS, fixed in 70% ice-cold ethanol at 4°C overnight, washed with PBS again, and stained with propidium iodide (200 µl of a 50 mg/L solution) at 37 °C for 20 min. Cell cycle profiles were assessed by measuring DNA content using flow cytometry.

### Immunofluorescence

Human TS cells maintained in the stem state or following differentiation were fixed with 4% paraformaldehyde (Sigma-Aldrich) for 20 min at room temperature. Cells were permeabilized with 0.1% Triton X-100. Immunofluorescence analysis was performed using a primary antibody against CYP1A1 (1:500, A3001; XenoTech) or AHR (1:500, MA1-514, Thermo Fisher). Alexa Fluor 488 goat anti-mouse IgG (1:800, A32723 Thermo Fisher), Alexa Fluor 568 goat anti-mouse IgG (1:800, A11031, Thermo Fisher), Alexa Fluor 568 goat anti-rabbit IgG, (1:800, A10042, Thermo Fisher) were used to detect locations of the primary antibody-antigen complexes within cells. Nuclei were visualized by staining with 4’6’-diamidino-2-phenylindole (**DAPI**, Molecular Probes). Images were captured on a Nikon 90i upright microscope with a Roper Photometrics CoolSNAP-ES monochrome camera.

### Short Hairpin RNA (shRNA) Constructs and Lentivirus Production

AHR shRNAs were subcloned into the pLKO.1 vector at *AgeI* and *EcoRI* restriction sites. shRNA sequences used in the analyses are provided in **Table S1**. Lentiviral packaging vectors were obtained from Addgene and included pMDLg/pRRE (plasmid 12251), pRSVRev (plasmid 12253), and pMD2.G (plasmid 12259). Lentiviral particles were produced following transient transfection of the shRNA-pLKO.1 vector and packaging plasmids into Lenti-X cells (632180, Takara Bio USA) using Attractene (301005, Qiagen) in Opti-MEM I (51985-034, Thermo Fisher). Cells were maintained in DMEM culture medium (11995-065, Thermo Fisher) supplemented with 10% FBS until 24 h prior to supernatant collection, at which time the cells were cultured in Basal TS Cell Medium supplemented with 200 μM L-ascorbic acid and 50 ng/mL of EGF.

### Lentiviral Transduction

Human TS cells were plated at 80,000 cells per well in 6-well tissue culture plates coated with 5 μg/mL collagen IV and incubated for 24 h. Immediately prior to transduction, medium was changed, and cells were incubated with 2.5 μg/mL polybrene for 30 min at 37°C.

Immediately following polybrene incubation, TS cells were transduced with 500 μL of lentiviral supernatant and then incubated for 24 h. Medium was changed at 24 h post-transduction and selected with puromycin dihydrochloride (5 μg/mL, A11138-03, Thermo Fisher) for two days. Surviving cells were cultured for one to three days in Complete Human TS Culture Medium before passaging and initiating EVT cell or ST differentiation.

### RNA Isolation, cDNA Synthesis, and Reverse Transcriptase-quantitative Polymerase Chain Reaction (RT-qPCR)

Total RNA was isolated from cells and tissues with TRIzol reagent (15596018, Thermo Fisher). cDNA was synthesized from 1 μg of total RNA using a High-Capacity cDNA Reverse Transcription kit (4368813; Thermo Fisher) and diluted 10 times with water. RT-qPCR was performed using a reaction mixture containing PowerSYBR Green PCR Master Mix (4367659; Thermo Fisher) and primers (250 nM each). PCR primer sequences are presented in **Table S2**. Amplification and fluorescence detection were carried out using a QuantStudio 7 Flex Real-Time PCR System (Thermo Fisher). An initial step (95 °C, 10 min) preceded by 40 cycles of a two-step PCR at: 92 °C, for 15 s and 60 °C for 1 min, followed by a dissociation step (95 °C for 15 s, 60 °C for 15 s, and 95 °C for 15 s). The comparative cycle threshold method was used for relative quantification of mRNA normalized to a housekeeping transcript, glyceraldehyde-3-phosphate dehydrogenase (*GAPDH*).

### RNA Sequencing (RNA-seq) Analysis

Transcript profiles were generated from human TS cells cultured in various differentiation states under control conditions or in the presence of AHR ligands (n=3/condition).

Complementary DNA libraries from total RNA samples were prepared with Illumina TruSeq RNA preparation kits (RS-122-2101, Illumina) according to the manufacturer’s instructions. RNA integrity was assessed using an Agilent 2100 Bioanalyzer (Agilent Technologies).

Barcoded cDNA libraries were multiplexed onto a TruSeq paired-end flow cell and sequenced (100-bp paired-end reads) with a TruSeq 200-cycle SBS kit (Illumina). Libraries were sequenced on Illumina HiSeq 2000 sequencer or Illumina NovaSeq 6000 at the University of Kansas Medical Center (**KUMC**) Genome Sequencing Facility. Reads from *.fastq files were mapped to the human reference genome (GRCh37) using CLC Genomics Workbench 12.0 (Qiagen).

Transcript abundance was expressed as reads per kilobase of transcript per million mapped reads (**RPKM**), and a false discovery rate of 0.05 was used as a cutoff for significant differential expression. Statistical significance was calculated by empirical analysis of digital gene expression followed by Bonferroni’s correction. Functional patterns of transcript expression were further analyzed using Ingenuity Pathway Analysis (Qiagen).

### Measurement of 2-Methoxyestradiol (2ME)

To assess 2ME biosynthesis TS cells were cultured in the presence of 17β-estradiol (10 nM). Conditioned medium from TS cells maintained in the stem state and following differentiation were collected and 2ME measured using an enzyme-linked immunosorbent assay (**ELISA**, 582261, Cayman Chemical).

### Western Blot Analysis

Cell lysates were prepared by sonication in radioimmunoprecipitation assay lysis buffer (sc-24948A, Santa Cruz Biotech) supplemented with Halt protease and a phosphatase inhibitor mixture (78443, Thermo Fisher). Protein concentrations were measured using the DC Protein Assay (5000113-115, Bio-Rad). Proteins (20 μg/lane) were separated by sodium dodecyl sulfate polyacrylamide gel electrophoresis and transferred onto polyvinylidene difluoride membranes (10600023, GE Healthcare). After transfer, membranes were blocked with 5% non-fat milk in Tris buffered saline with 0.1% Tween 20 (**TBST**) and probed with primary antibodies to AHR (1:1000 dilution, MA1-514, Thermo Fisher) or GAPDH (1:1000 dilution, ab9485, Abcam) overnight at 4°C. Membranes were washed three times for five min with TBST and then incubated with secondary antibodies (goat anti-rabbit IgG HRP, A0545; Sigma-Aldrich and goat anti-mouse IgG HRP, 7076; Cell Signaling) for 1 h at room temperature. Immunoreactive proteins were visualized by enhanced chemiluminescence (Amersham).

### Statistical Analysis

Statistical analyses were performed with GraphPad Prism 9 software. Welch’s *t* tests, Brown–Forsythe and Welch analysis of variance (ANOVA) and Two-way ANOVA were applied as appropriate (see figure legends for additional information). Statistical significance was determined as P<0.05.

## RESULTS

### Examination of the effects of AHR activation on human trophoblast cells

We examined the effects of AHR ligand TCDD in human TS cells at three developmental states: i) stem cell state, ii) EVT cell differentiation state, and iii) ST differentiation state.

#### Stem cell state

Human TS cells can expand and exhibit a signature transcript profile when maintained in a condition to promote the stem cell state (**Okae et al. 2018; Varberg et al. 2023**). *CYP1A1* and *CYP1B1* increased dramatically in response to TCDD exposure (**Figure 1A-C**). Exposure of TS cells maintained in the stem state to TCDD (10 nM) did not adversely affect cell viability or cell cycle (**Figure S1**). Analysis of RNA-seq of TCDD treated versus control cells resulted in the identification of 668 differentially expressed genes (**DEGs**), including 484 genes upregulated and 184 genes downregulated by exposure to TCDD (10 nM) (**Figure 1D and Dataset 1**). This differential gene expression pattern was validated by RT-qPCR (**Figure 1E** and **F**). Functional pathways affected by TCDD exposure, included pathways associated with protein translation and cell-extracellular matrix adhesion (**Figure S2**). There was some indication that TCDD exposure to TS cells in the stem state enhanced expression of ST-associated transcripts (e.g. *CGB5, CGB7, LEP, SLC7A5*; **Dataset 1**). We also examined the consequences of human TS cell TCDD exposure (10 or 25 nM) during the stem state (24 h) on subsequent EVT cell and ST differentiation. TCDD exposure during the stem state did not adversely affect EVT cell or ST differentiation (**Figure S3**).

**Figure 1.**
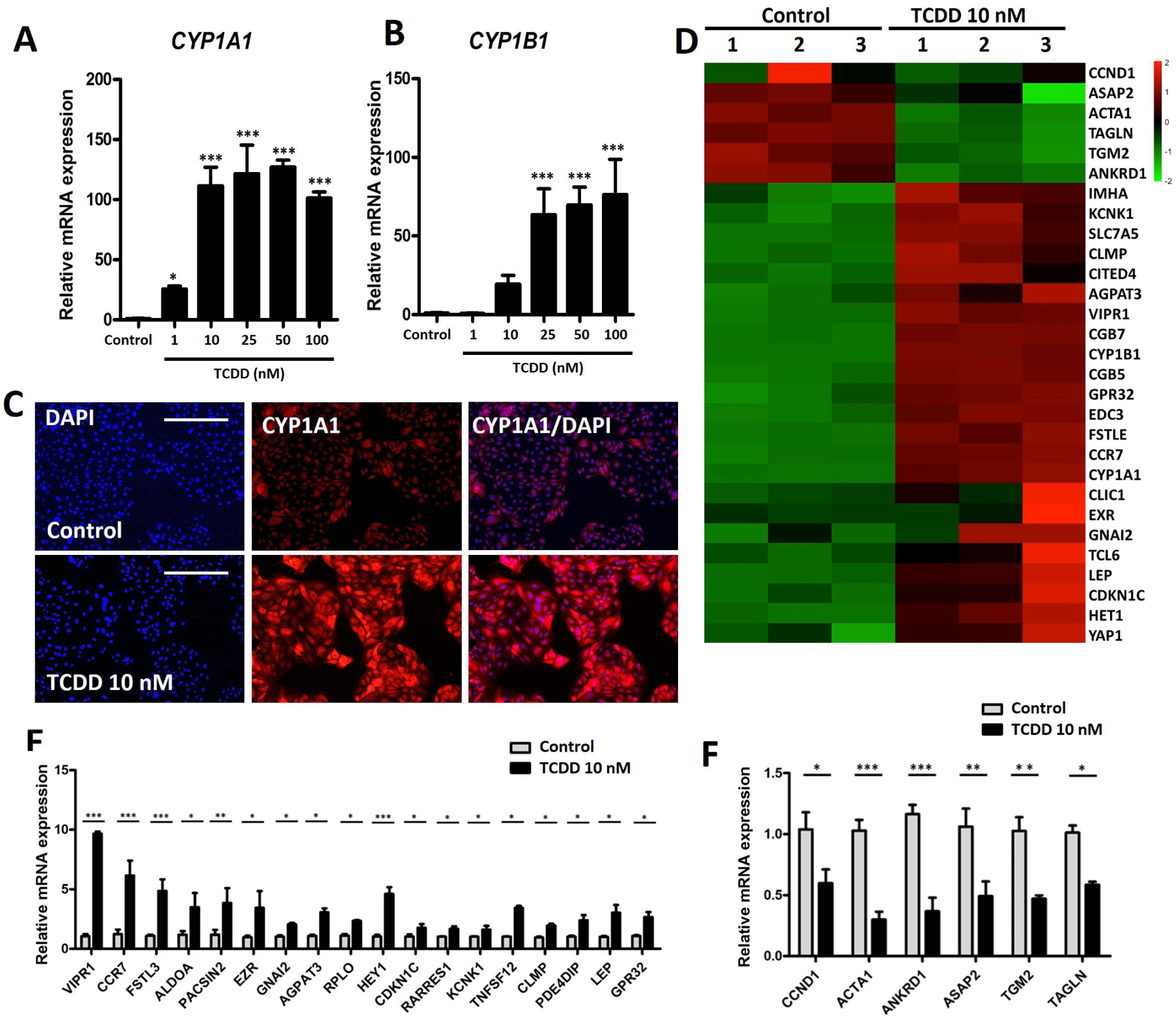
Effects of AHR activation on human TS cells in the stem state. (A, B) *CYP1A1* and *CYP1B1* transcript levels in human TS cells exposed to Control conditions or TCDD (1-100 nM) for 24 h. (C) Immunocytochemistry of CYP1A1 protein expression in human TS cells exposed to Control conditions or TCDD (10 nM) for 24 h (Scale bar: 300 μm). DAPI identifies cell nuclei (blue). (D) Heatmap showing select transcripts from RNA-seq analysis of human TS cells exposed to Control conditions or TCDD (10 nM) 24 h. (E, F) RT-qPCR validation of selected up regulated and down-regulated transcripts in human TS cells exposed to Control or TCDD (10 nM). n=3. Graphs represent mean values ± standard error of the mean (**SEM**), unpaired t test, *P < 0.05, **P < 0.01, and ***P < 0.001.

#### EVT cell differentiation state

TCDD exposure did not adversely affect the morphology of differentiated EVT cells (**Figure 2A**); however, TCDD exposure did increase *CYP1A1* and *CYP1B1* transcript levels and CYP1A1 protein expression (**Figure 2B** and **C**). RNA-seq analysis of control and TCDD exposed cells identified 336 DEGs, including 173 upregulated transcripts and 163 downregulated transcripts in TCDD treated cells (**Dataset 2, Figure 2D**). RT-qPCR validation of a subset of these transcripts is shown (**Figure 2E** and **F**). Functional pathways affected by TCDD exposure, included pathways associated with protein translation, cell-cell interactions, and cell death (**Figure S4**).

**Figure 2.**
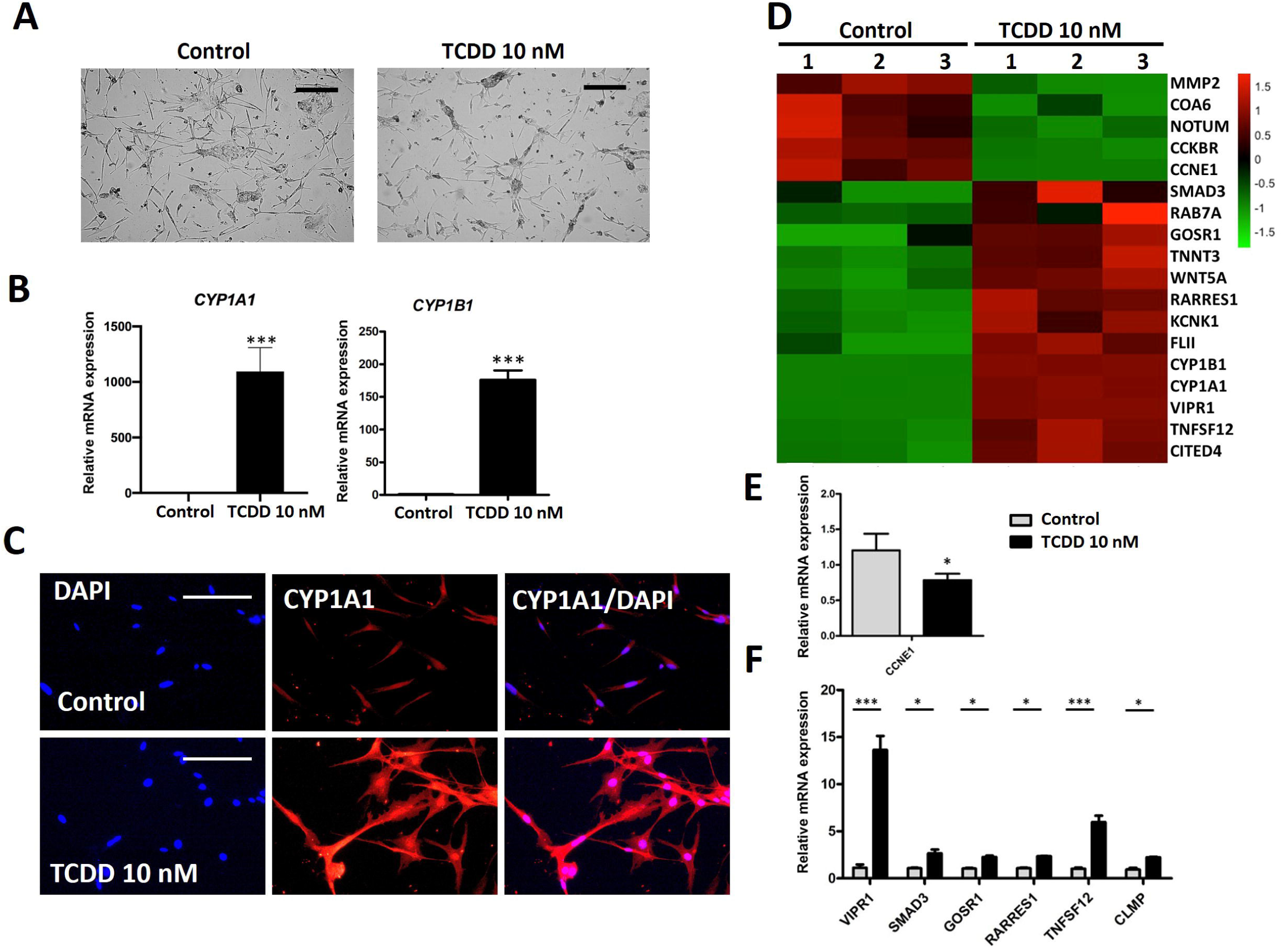
Effect of AHR activation in EVT cells. (A) Phase-contrast images depicting cell morphology of EVT cells differentiated from human TS cells in presence of vehicle or TCDD (10 nM) (Scale bar = 500 μm). (B) Expression of *CYP1A1* and *CYP1B1* following exposure to vehicle or TCDD (10 nM) during EVT cell differentiation. (C) Immunofluorescence of CYP1A1 expression (red) in EVT cells treated with vehicle and TCDD (10 nM) (Scale bar: 300 μm). DAPI marks cell nuclei (blue). (D) Heatmap showing select transcripts from RNA-seq analysis of EVT cells exposed to vehicle versus TCDD (10 nM). (E, F) RT-qPCR validation of selected up-regulated and down-regulated transcripts in vehicle versus TCDD treated cells. n = 3. Graphs represent mean values ± SEM, unpaired t test, *P < 0.05, **P < 0.01, and ***P < 0.001.

#### ST differentiation state

TCDD exposure did not adversely affect the morphology of differentiated ST (**Figure 3A**); however, TCDD exposure during ST differentiation did increase the expression of *CYP1A1* and *CYP1B1* (**Figure 3B**). RNA-seq analysis of control and TCDD exposed cells identified 353 DEGs, including 154 upregulated genes and 199 downregulated genes in TCDD treated cells (**Dataset 3, Figure 3C**). RT-qPCR validation of a subset of these transcripts is shown (**Figure 3D**). Functional pathways affected by TCDD exposure, included pathways associated with estrogen biosynthesis and AHR and hypoxia signaling (**Figure S5**).

**Figure 3.**
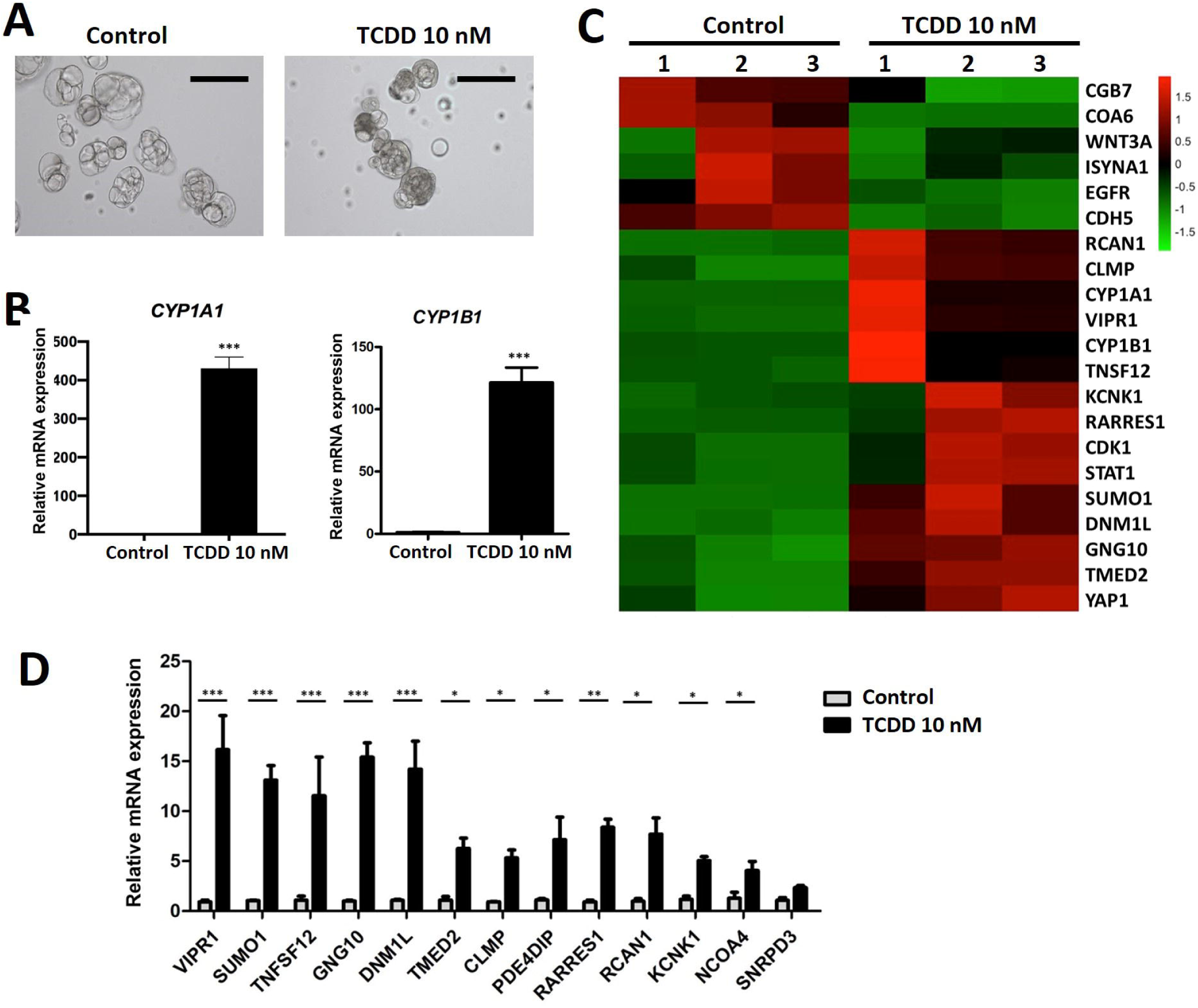
Effect of AHR activation in syncytiotrophoblast differentiation. (A) Phase-contrast images depicting three-dimensional (**3D**) syncytiotrophoblast development in presence of vehicle or TCDD (10 nM) (Scale bar = 300 μm). (B) Expression of *CYP1A1* and *CYP1B1* following exposure to vehicle or TCDD (10 nM) during 2D and 3D syncytiotrophoblast differentiation. (C) Heatmap showing select transcripts from RNA-seq analysis of 3D syncytiotrophoblast exposed to vehicle versus TCDD (10 nM). (D) RT-qPCR validation of selected up-regulated transcripts in cells treated with vehicle versus TCDD, n=3. Graphs represent mean values ± SEM, unpaired t test, *P < 0.05, **P < 0.01, and ***P < 0.001.

Cells in each trophoblast cell differentiation state exhibited similar TCDD induced activation of *CYP1A1* and *CYP1B1* (**Figures 1-3**). However, based on the total number of DEGs, TS cells in the stem state were maximally responsive to TCDD (668 DEGs), whereas EVT cells were the least responsive to TCDD (336 DEGs). ST exhibited an intermediate response to TCDD (353 DEGs). These observations indicate that TS cells in the stem state may be more vulnerable to TCDD exposure than differentiated trophoblast cells. Genes associated with differentially regulated pathways are presented in **Dataset 4**. The molecular basis for the increased TCDD sensitivity of TS cells in the stem state is not known.

#### Role of AHR in TCDD induction of *CYP1A1* and *CYP1B1*

We next tested whether TCDD effects on *CYP1A1* and *CYP1B1* expression in human TS cells were dependent upon AHR using a loss-of-function approach. AHR expression was silenced in human TS cells using stable lentiviral-mediated delivery of control and AHR-targeted shRNAs. Disruption of AHR expression was verified by RT-qPCR, western blotting, and immunofluorescence (**Figure 4A-C**). AHR knockdown TS cells maintained in the stem state did not effectively respond to TCDD with an induction of *CYP1A1* and *CYP1B1* expression (**Figure 4D**). The results demonstrated that TCDD induction of *CYP1A1* and *CYP1B1* gene expression is AHR dependent.

**Figure 4.**
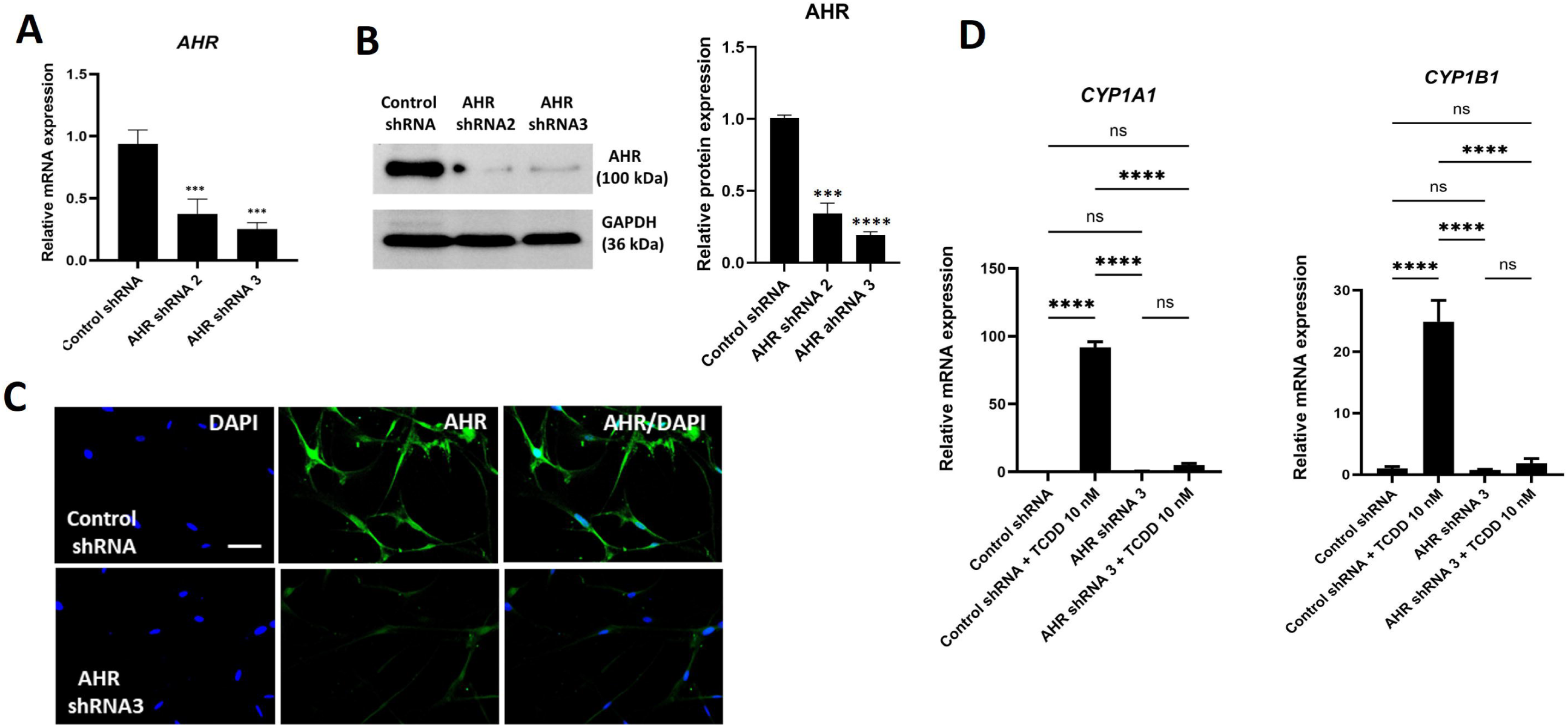
AHR dependent activation of CYP1A1 in human TS cells. RT-qPCR (A) and western blot (B) assessment of lentiviral vector-mediated AHR silencing efficiency in human TS cells expressing control or AHR shRNAs. (C) Immunocytochemistry of AHR protein expression (green) in control shRNA or AHR shRNA silenced cells (Scale bar: 300μm). DAPI marks cell nuclei (blue). (D) *CYP1A1* and *CYP1B1* transcript level measurements in control shRNA or AHR shRNA silenced cells in presence of TCDD (10 nM) for 24 h. n=3. Graphs represent mean values ± SEM, one-way ANOVA analysis, Tukey’s post hoc test. *P < 0.05, **P < 0.01, ***P < 0.001, ****P < 0.0001.

#### AHR activation drives 2ME biosynthesis in human TS cells

In the above experimental results presented in Figures 1, 2, and 3, we observed significant effects of TCDD exposure on gene expression but not on the maintenance of the human TS cell stem state or in the capacity for human TS cells to differentiate into EVT cells or ST. CYP1A1 expression was especially responsive to TCDD and has the capacity to transform endogenous and exogenous compounds, including 17β estradiol, into biologically active molecules such as 2ME (**Thomas and Potter 2013**). Consequently, we examined the effects of TCDD on the capacity of human TS cells in the presence of 17β estradiol to synthesize 2ME. TCDD exposure significantly stimulated 2ME biosynthesis in human TS cells maintained in the stem cell state (**Figure 5A**) and TS cells induced to differentiate into EVT cells or ST (**Figure 5B**). A pathway showing the involvement of AHR, CYP1A1, and 2ME in xenobiotic action at the placentation site is shown (**Figure 5C**).

**Figure 5.**
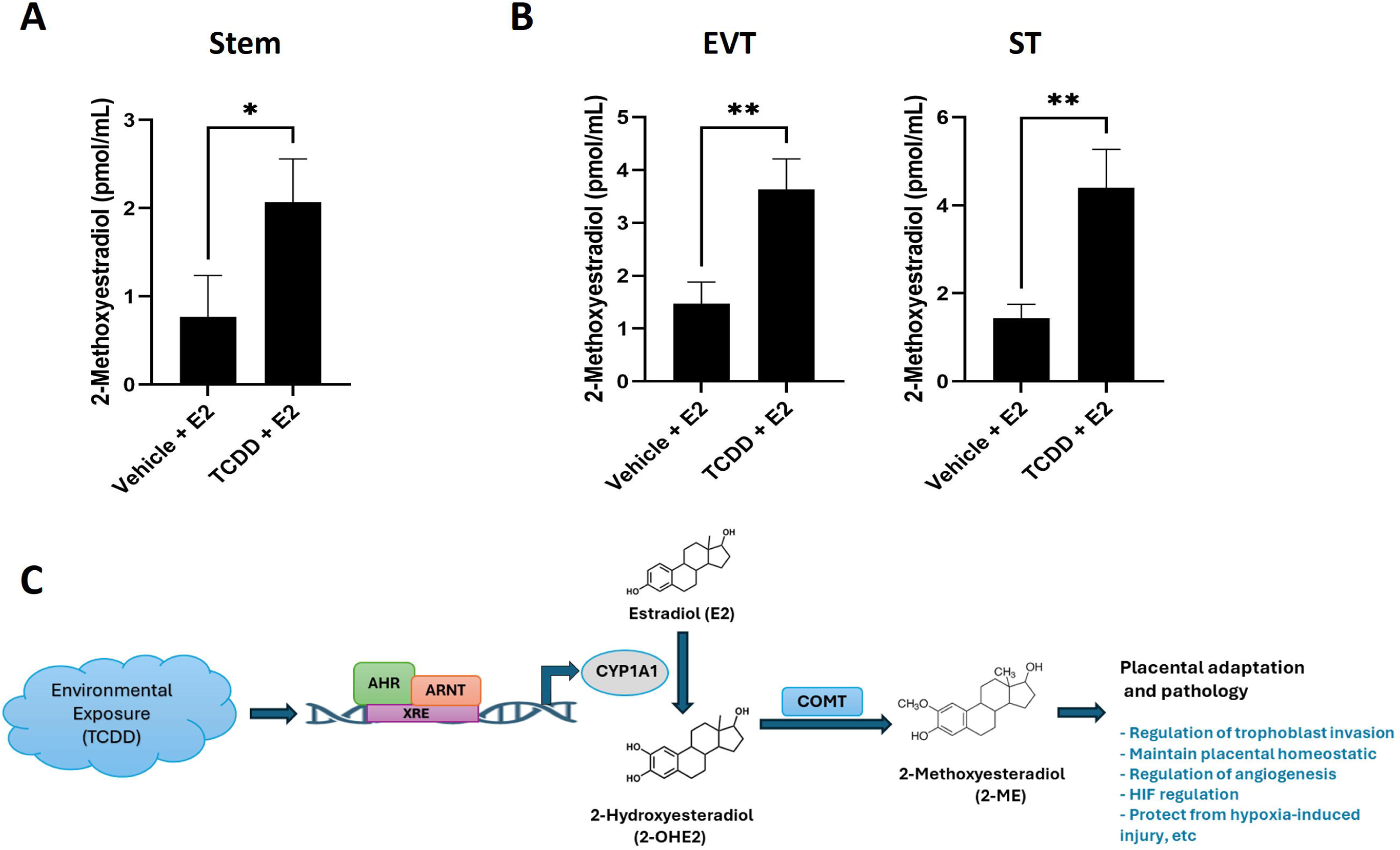
Effects of AHR activation on 2-methoxyestradiol production by human TS cells. 2-methoxyestradiol concentration (pmole/mL) measured in TS cells maintained in the stem state (A) or induced to differentiate into EVT cells or ST (B). Cells were exposed to vehicle + 17β-estradiol (E2; 10 nM) or TCDD (10 nM) + E2 (10 nM) for 48 h before harvesting conditioned medium for 2-methoxyestradiol measurement. n=3. Graphs represent mean values ± SEM, unpaired t test, *P < 0.05, and **P < 0.01 (C) Schematic of a TCDD-mediated pathway affecting placentation. Abbreviations: AHR, aryl hydrocarbon receptor; ARNT, aryl hydrocarbon receptor nuclear translocator; XRE, xenobiotic response element; COMT, catechol-O-methyltransferase.

## DISCUSSION

The primary interface between mother and fetus, the placenta, serves two critical functions: extraction of nutrients from the maternal compartment and facilitation of nutrient delivery to the developing fetus. This delivery system also serves as a barrier to environmental exposures. The aryl hydrocarbon receptor (**AHR**) is an important component of the barrier. AHR signaling is activated by environmental pollutants and toxicants that can potentially affect cellular and molecular processes, including those controlling trophoblast cell development and function. In this report, we discovered that TCDD activates AHR signaling in human trophoblast cells and evokes a robust transcriptional response, which includes stimulating CYP1A1 expression. Human trophoblast cells have similarly been shown to respond to AHR ligands with an increase in CYP1A1 gene expression (**Stejskalova et al. 2011; Wakx et al. 2018).** These TCDD activated changes in TS cells did not adversely affect their ability to self-renew or to differentiate into either EVT cells or ST. However, they can affect the availability of biologically active ligands at the maternal-fetal interface.

TCDD does not act directly to adversely affect the development of rat or human trophoblast cells (**Iqbal et al. 2021; present study**). Rat trophoblast cells lack the requisite cellular machinery needed to respond to TCDD (**Iqbal et al. 2021**), while human trophoblast cells are responsive to TCDD, but without negative consequences on TS cell self-renewal, maintenance of the TS cell stem state or the differentiation of TS cells into EVT cells and ST (**present study**).

Although, trophoblast cell development is not adversely affected by direct exposure to TCDD in the rat or human, there are pronounced consequences of AHR activation on placentation and trophoblast cell function. In the rat, TCDD exposure alters placentation and the behavior of trophoblast cells via indirect actions on endothelial cells (**Iqbal et al. 2021**). Since human endothelial cells are also responsive to AHR ligands (**Bredhult et al. 2007; Kopf and Walker 2010; Li et al. 2015**), we expect similar endothelial cell-mediated in vivo actions of TCDD on human placentation. In addition, as shown here, TCDD-mediated activation of AHR signaling also possesses the capacity to directly fine-tune trophoblast cell function in each of its developmental states.

The relevance of species differences in trophoblast cell engagement with its environment is unknown. At one level, survival of a species would appear to be enhanced by the ability to actively adapt to the environment, especially through the upregulation of an enzyme that can transform a potentially dangerous compound into a compound that can be made less threatening or extricated from the body. This implies that the actions of environmentally activated enzymes possessing biotransformational properties are unilateral in their efforts. This is not the case for AHR and its downstream targets, especially CYP1A1. Endogenous AHR ligands are present in the cellular milieu (**Nguyen and Bradfield, 2008**) and CYP1A1 can act on endogenous compounds (**Stejskalova and Pavek, 2011; Bock 2014**). Among the endogenous compounds that CYP1A1 can act on is the steroid hormone, 17β estradiol (**Thomas and Potter 2013**).

Biosynthesis of estrogens represents a key species difference in the evolution of the placenta (**Soares et al. 2018**). Trophoblast cells of the human placenta possess aromatase (cytochrome P450 family 19 subfamily A member 1, CYP19A1), the enzyme responsible for conversion of androgens to estrogens (**Albrecht and Pepe 1990; Simpson et al. 1997**), whereas this key enzyme in estrogen biosynthesis is not present in placentas of the rat and mouse (**Kamat et al. 2002**). Interestingly, estrogen action and AHR signaling have been linked (**Tarnow et al. 2019**). Thus, species differences in trophoblast cell responses to environmental signals capable of activating AHR signaling may be linked to species differences in placental capacity for estrogen biosynthesis. Investigating the relationship of AHR signaling and estrogen biosynthesis in placentas of other species could be informative.

The most prominent effect of AHR activation on human trophoblast cells was on the expression of CYP1A1. CYP1A1 does little to affect cell function unless there is a substrate for it to act on. As indicated above, 17β estradiol is a notable CYP1A1 substrate produced within the human placenta. Estrogens are prominent activators of two nuclear estrogen receptors (**Deroo and Korach 2006**), which are critical for reproductive function, including the establishment and maintenance of pregnancy (**Deroo and Korach 2006; Hewitt et al. 2016**). CYP1A1 can hydroxylate estradiol to 2-hydroxyestradiol and 4-hydroxyestradiol (catechol estrogens) (**Thomas and Potter 2013; Kumar et al. 2016**). These modifications of estradiol decrease its availability for signaling through nuclear estrogen receptors and generate biologically active compounds with different properties. Catechol-O-methyltransferase (**COMT**) can modify 2-hydroxyestradiol to 2ME (**Thomas and Potter 2013; Kumar et al. 2016**). Human trophoblast cells exposed to TCDD exhibit an enhanced capacity to convert estradiol to 2ME (**present study**). 2ME is a compound with biological functions implicated in regulatory processes associated with angiogenesis, cellular responses to hypoxia, and preeclampsia (**Mabjeesh et al. 2003; Kanasaki et al. 2008; Lee et al. 2010; Perez-Sepulveda et al. 2013; Pinto et al. 2014**).

Collectively, these findings indicate that TCDD, a prototypical AHR ligand, has the capacity to influence the behavior of trophoblast cells within the human maternal-fetal interface and potentially pregnancy outcomes.

## ACKNOWLEDGMENTS

We thank Stacy Oxley, Leslie Tracy, and Brandi Miller for their assistance. Research was supported by KUMC BRTP and K-INBRE P20 GM103418 (VS), National Institutes of Health: ES028957 (KI) and HD020676, ES029280, HD105734 and the Sosland Foundation.

## DATA SHARING

The datasets generated and analyzed for this study have been deposited in the Gene Expression Omnibus (**GEO**) database, https://www.ncbi.nlm.nih.gov/geo/ (accession no. GSE246513).

## SUPPLEMENTARY FIGURES

**Figure S1.** Effects of TCDD on human cell cycle and cell death. (A, B) Human TS cells were stained with annexin-V (**AV**) and propidium iodide (**PI**) and subjected to flow cytometry to determine cell death. Human TS cells were treated with vehicle or TCDD (10 and 100 nM). (C, D) Human TS cells were stained with PI and subjected to flow cytometry to determine DNA content and stage of the cell cycle. n=2. Graphs represent mean values ± SEM, Two-way ANOVA, *P < 0.05, ***P < 0.001, and ****P < 0.0001.

**Figure S2.** Pathway analysis of RNA-sequencing datasets of human TS cells maintained in the stem state exposed to vehicle or TCDD (10 nM).

**Figure S3.** Effects of TCDD exposure in the stem state on the capacity of human TS cells to differentiate. Human TS cells were treated in the stem state with TCDD (10 nM) for 24 h and then induced to differentiate into EVT cells or ST. (A) Morphology of human TS cells induced to differentiated into EVT cells. (B) RT-qPCR measurement of *HLA-G* and *MMP2* levels, transcripts associated with EVT cell differentiation. (C) Morphology of human TS cells induced to differentiated into ST. (D) RT-qPCR measurement of *CGB5* and *SDC1* levels, transcripts associated with ST differentiation.

**Figure S4.** Pathway analysis of RNA-sequencing datasets of human TS cells induced to differentiate into EVT cells exposed to vehicle or TCDD (10 nM).

**Figure S5.** Pathway analysis of RNA-sequencing datasets of human TS cells induced to differentiate into ST exposed to vehicle or TCDD (10 nM).

