## Supplemental File for "Aryl Hydrocarbon Receptor Activation Drives 2-Methoxy Estradiol Secretion in Human Trophoblast Stem Cell Development"

**SUPPLEMENTARY FIGURES**

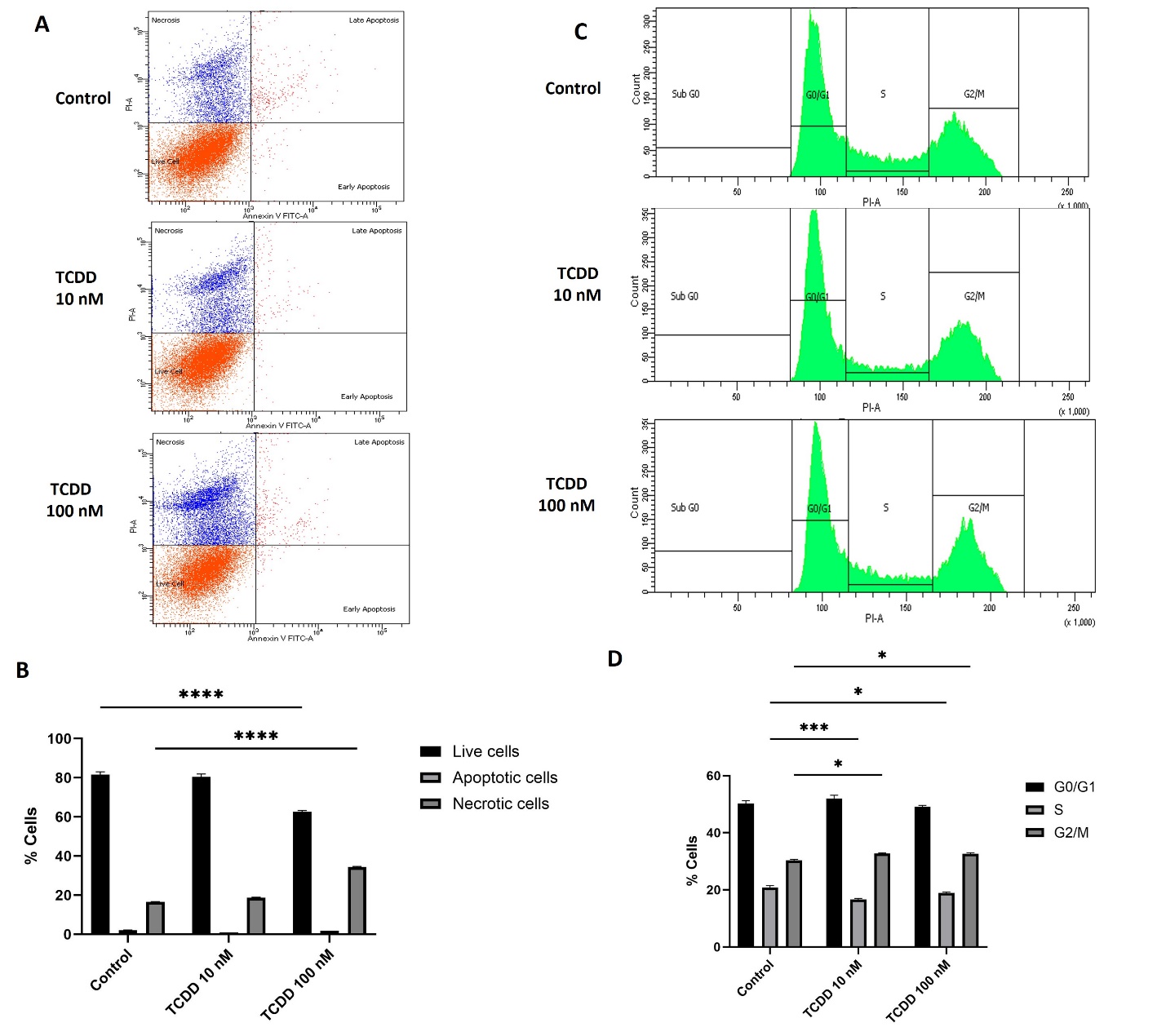

**Figure S1.** Effects of TCDD on human cell cycle and cell death. (A, B) Human TS cells were stained with annexin-V (**AV**) and propidium iodide (**PI**) and subjected to flow cytometry to determine cell death. Human TS cells were treated with vehicle or TCDD (10 and 100 nM). (C, D) Human TS cells were stained with PI and subjected to flow cytometry to determine DNA content and stage of the cell cycle. n=2. Graphs represent mean values ± SEM, Two-way ANOVA, *P < 0.05, ***P < 0.001, and ****P < 0.0001.

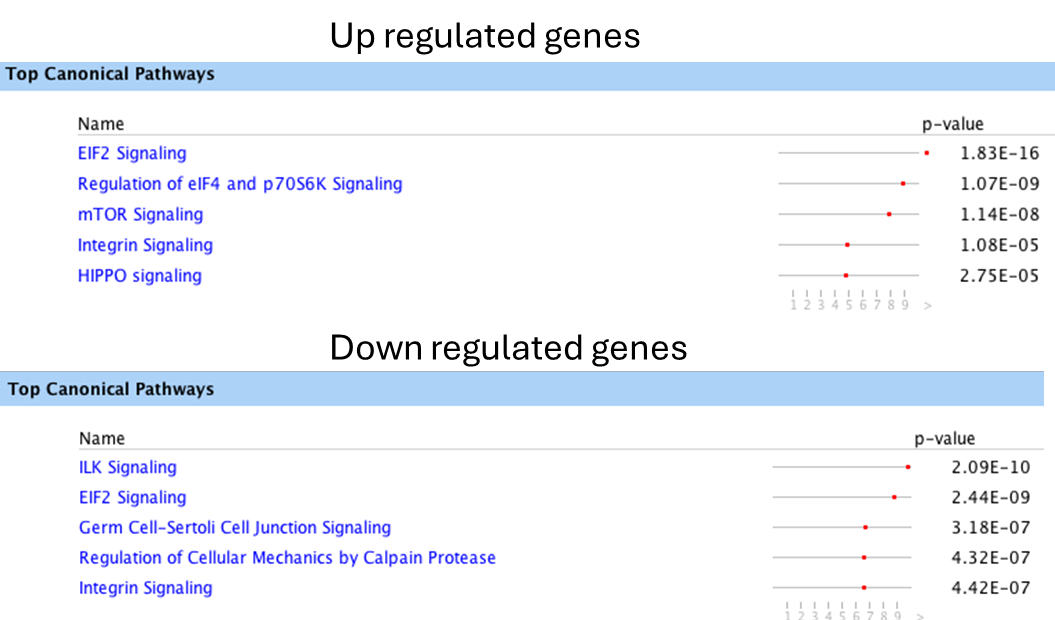

**Figure S2.** Pathway analysis of RNA-sequencing datasets of human TS cells maintained in the stem state exposed to vehicle or TCDD (10 nM).

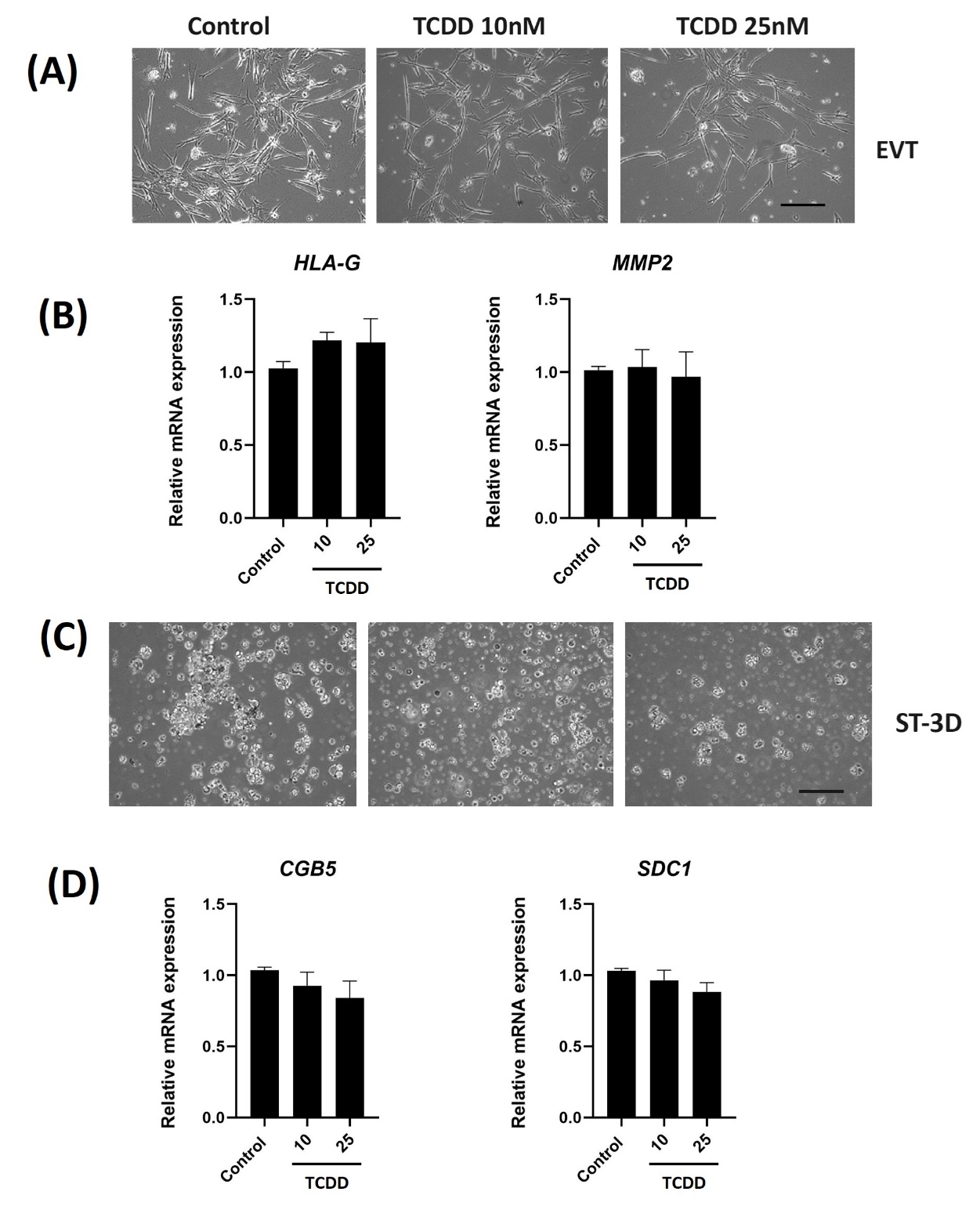

**Figure S3.** Effects of TCDD exposure in the stem state on the capacity of human TS cells to differentiate. Human TS cells were treated in the stem state with TCDD (10 nM) for 24 h and then induced to differentiate into EVT cells or ST. (A) Morphology of human TS cells induced to differentiated into EVT cells. (B) RT-qPCR measurement of *HLA-G* and *MMP2* levels, transcripts associated with EVT cell differentiation. (C) Morphology of human TS cells induced to differentiated into ST. (D) RT-qPCR measurement of *CGB5* and *SDC1* levels, transcripts associated with ST differentiation.

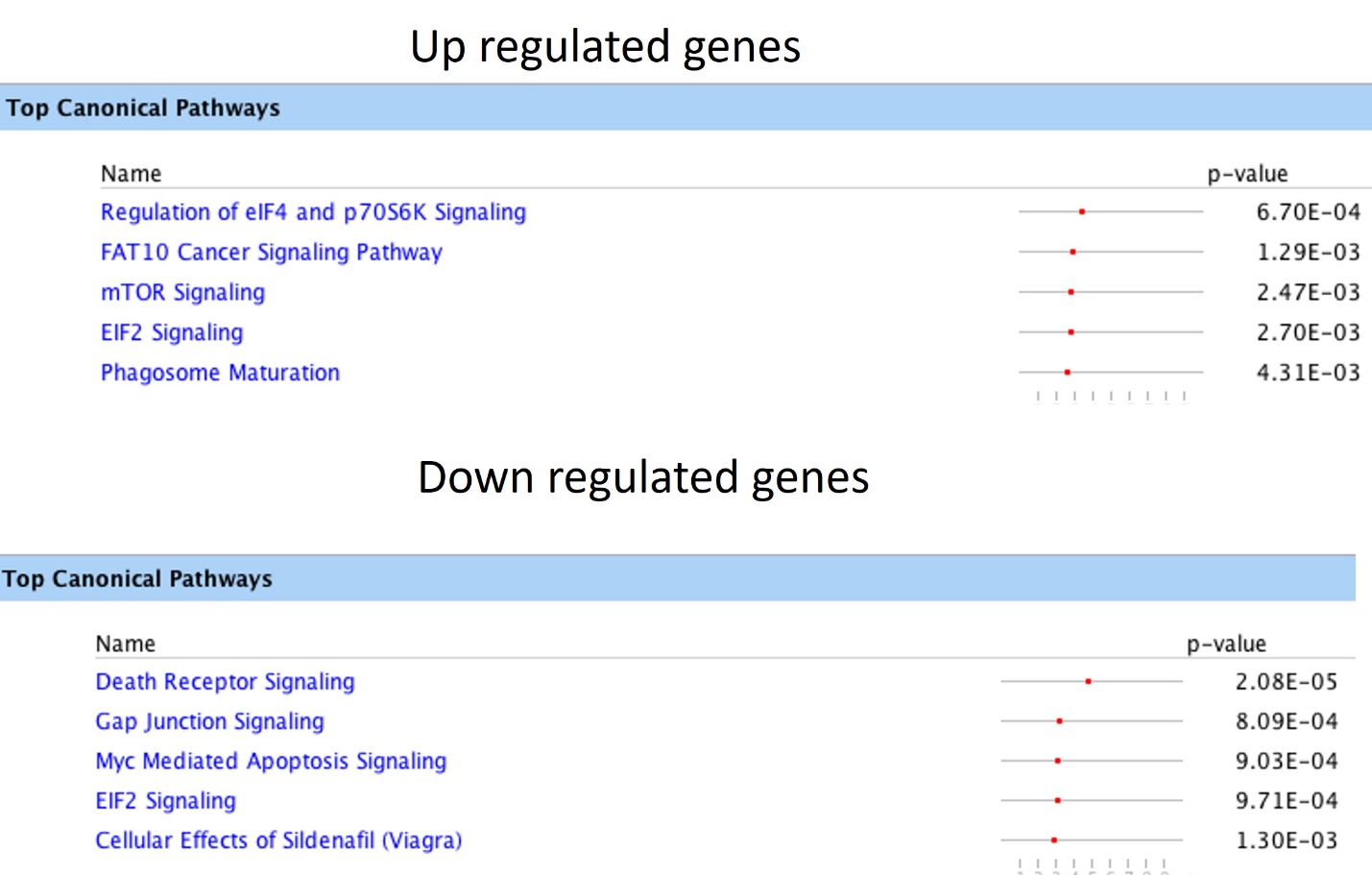

**Figure S4.** Pathway analysis of RNA-sequencing datasets of human TS cells induced to differentiate into EVT cells exposed to vehicle or TCDD (10 nM).

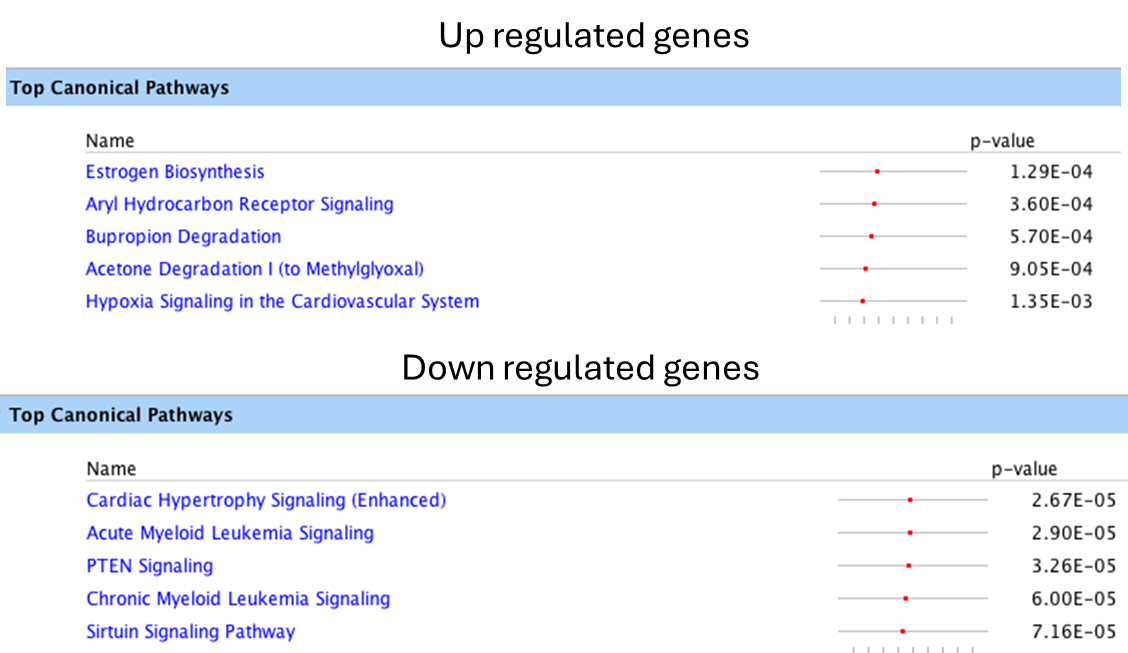

**Figure S5.** Pathway analysis of RNA-sequencing datasets of human TS cells induced to differentiate into ST exposed to vehicle or TCDD (10 nM).

**Table S1. shRNA sequences**

| **shRNA** | **Sequences** |
| --- | --- |
| AHR shRNA 2 | ACTGCTTAAAGTTGGTATTAA |
| AHR shRNA 3 | GCAACAAGATGAGTCTATTTA |

**Table S2. Primer sequences for qRT-PCR analyses**

| **Target** | **Species** | **Forward** | **Reverse** |
| --- | --- | --- | --- |
| *CYP1A1* | Human | ACC TTC CCT GAT CCT TGT GA | GGA GAT TGG GAA AAG CAT GA |
| *CYP1B1* | Human | TCA ACA AGG ACC TGA CCA GC | TCA TTT GGG TTG GCC CTG AA |
| *VIPR1* | Human | GAA CTC TGT GTG GTG CGT CT | AGT TTC CTC ATC TCT GCC GTC |
| *CCR7* | Human | GAC ACA GGC ATA CCT GGA AA | TGG TTT TAC CGC CCA GAG AG |
| *FSTL3* | Human | CTA CAT CTC CTC GTG CCA CA | TCT TCT GCA GAC TCA CCA CCT |
| *ALDOA* | Human | GGT AGT AGC AAG TTC CTG GCA | CCC GCG TTC TCT CCT TGA A |
| *PACSIN2* | Human | GGG ACA TTA AGG GTG CTG CT | TCC GCA GAC CTG AAT CGA AC |
| *EZR* | Human | ACT CGG ACA TTG ATT GGT TTC G | CGG GCG CTC TAA GGG TTC T |
| *GNAI2* | Human | TCC TCG TGG ATG ATC TTC ATC TG | CGG CCG AGC GCT CTA AG |
| *AGPAT3* | Human | AGC ACG AAC TGG GTC TTC AG | CGG CTG CAG GAC GGC |
| *RPLPO* | Human | AGA CGA TGT CAC TTC CAC GAG | GCG GTT TCT GAT TGG CTA CTT TGT |
| *HEY1* | Human | TCC TGC CGT ATG CAG CAT TT | GCT TTT GAG AAG CAG GGA TCT |
| *CDKN1C* | Human | GCG GCG ATC AAG AAG CTG T | GCT TGG CGA AGA AAT CGG AGA |
| *RARRES1* | Human | ACT AGT GTG AGG CAG TGG GT | GCA TGA ATT CAG TCT AAG GAG ACC |
| *KCNK1* | Human | CTT GCT CTA CCT GGT CTT CGG | TGT GGC CAT AAC CTG TGG T |
| *TNFSF12* | Human | CCT CGC AGA AGT GCA CCT AAA | TCA GGT AGA CAG CCT TCC CC |
| *CLMP* | Human | TTA CTG GCA GCG AAT CCG AG | CAG GGT GGT TGT AGT CAA TCC |
| *PDE4DIP* | Human | CCT GGC AAA GTG GGA GAA TC | GGC TGG AGT ACA TGG CAG A |
| *LEP* | Human | AAT GCA TTG GGG AAC CCT GT | AGG AGA CTG ACT GCG TGT GT |
| *GPR32* | Human | TGG ACC GTT GCA TCT CTG TC | GAA TTT CAG GTG CGC AGA GC |
| *CCND1* | Human | CCA GGT TCC ACT TGA GCT TGT T | CTG TGC ATC TAC ACC GAC AAC TC |
| *ACTA1* | Human | GCG GGG CGA TGA TCT TGA | CAC GAT GTA CCC TGG GAT CG |
| *ANKRD1* | Human | GTC TGC CTC ACA GGC GAT AA | AGC GCC CGA GAT AAG TTG C |
| *ASAP2* | Human | CAT TTT CCA CGT GAG CCA GC | CCC ATG AGG ACT ACA AGG CG |
| *TGM2* | Human | GTA CAC AGC ATC CGC GGT C | CAC TTT GAG GGC CGC AAC TA |
| *TAGLN* | Human | CTG AGG AAG CCT TCT TTC CCC | GGG CCA CAC TGC ACT ATG AT |
| *CCNE1* | Human | CAG CCC CAT CAT GCC GAG | TAT TGT CCC AAG GCT GGC TC |
| *SMAD3* | Human | AAG CGC ACT GAC CAT AAG AG | CCA TCC AGG GAC TCA AAC G |
| *GOSR1* | Human | GTA CCC GAG ATG GAA GAC GC | TTA CCC CTG TAA GCC TTG CC |
| *SUMO1* | Human | TGT GGG GAA GGG AGA AGG AT | AAG GTT TTG CCT CCT GGT CA |
| *GNG10* | Human | CGT GGA GAG GAT CAA GGT CTC | GAG TCT TCA GAG TAA AGC ACA GG |
| *DNM1L* | Human | TCA CCC GGA GAC CTC TCA TT | TCT GCT TCC ACC CCA TTT TCT |
| *TMED2* | Human | GAG ATA CCT GGG ATG CTC AC | TTA CAG CTG TCA TCG CCA CT |
| *RCAN1* | Human | GAG GCG GCC AGA TGA ACT C | GGG ACT CAA ATT TGG CCT TGC T |
| *NCOA4* | Human | GTC TTG CAT GAC TGG CCT AAG | TCT CCT CAC TGC TCC TGT ATC A |
| *SNRPD3* | Human | AGG CCA GAG CCG AAC TCT C | CAC CGG TGT TCG TCT CAC AT |
| *AHR* | Human | TAC AGA GTT GGA CCG TTT GG | GCC TCC GTT TCT TTC AGT AG |
| *GAPDH* | Human | CCTCAACGACCACTTTGTCAAG | TCTTCCTCTTGTGCTCTTGCTG |

**Dataset 1 (separate file).** Differentially regulated transcripts identified by RNA-seq analysis in STEM human X,X CT-27 TS cells following TCDD exposure.

**Dataset 2 (separate file).** Differentially regulated transcripts identified by RNA-seq analysis in day 8 EVT differentiated human X,X CT-27 TS cells following TCDD exposure.

**Dataset 3 (separate file).** Differentially regulated transcripts identified by RNA-seq analysis in day 6 ST differentiated human X,X CT-27 TS cells following TCDD exposure.

**Dataset 4 (separate file).** Genes associated with differentially top canonical pathways.
